# Endothelial MRG15 Is a Mechanosensitive Suppressor of Atherosclerosis

**DOI:** 10.64898/2026.01.28.702256

**Authors:** Yongbo Li, Xiaoke Lin, Jinghua Ma, Qingjun Jiang, Jiayi Lin, Lefeng Qu, Qiurong Ding, Xiangxiang Wei, Xiuling Zhi, Panpan Zhang, John Y.-J. Shyy, Elena Osto, Yuxiang Dai, Jieyu Guo, Dan Meng

## Abstract

Disturbed blood flow induces early endothelial inflammation in atherosclerosis, yet the precise mechanisms of endothelial sensing of disturbed flow remain incompletely understood. In this study, we integrated proteomic profiling of human endothelial cells (ECs) subjected to disturbed flow with coronary artery disease risk related genes from large-scale genome-wide association studies (GWAS). We identified mortality factor 4-like protein 1 (MORF4L1, also called MRG15) as a critical modulator in the pathogenesis of atherosclerosis induced by disturbed flow. We show that the expression of MRG15 is markedly reduced in ECs of human aortic atherosclerotic plaques, as well as in ECs exposed to disturbed flow in mouse carotid artery and in cultured human umbilical vein endothelial cells (HUVECs). Endothelial-specific deletion of *Mrg15* significantly worsened, while its overexpression attenuated endothelial inflammation and atherosclerotic lesions in turbulent blood flow- or Western diet-induced mouse atherosclerosis model. Single-cell transcriptomics showed that *Mrg15* deficiency increased endothelial inflammation and intercellular adhesion molecule 1 (*Icam1*) expression, and enhanced integrin-mediated adhesion pathways. Mechanistically, MRG15 facilitated the recruitment of enhancer of zeste homolog 2 (EZH2) to maintain repressive histone H3 lysine 27 trimethylation (H3K27me3) marks on the promoters of *ICAM1* and integrin subunit alpha 5 (*ITGA5*). Disturbed blood flow rapidly led to an elevation of protein neddylation, which subsequently induced neddylation-dependent degradation of MRG15 within endothelial cells. This degradation of MRG15 alleviated the transcriptional repression of *ICAM1* and *ITGA5*, thereby enhancing monocyte adhesion to ECs. These findings highlight endothelial MRG15 as a mechanosensitive suppressor of atherosclerosis induced by disturbed flow. Consequently, MRG15 emerges as a promising novel therapeutic target for atherosclerosis.

## Introduction

Atherosclerosis remains the primary cause of global cardiovascular mortality(*1*). Lesions predictably develop at arterial branches and curvatures—sites exposed to oscillatory shear stress (OSS), while straight segments subjected to unidirectional laminar shear stress (LSS) are largely protected(*2, 3*). Vascular endothelial cells (ECs) form the primary interface between blood flow and the vessel wall, functioning as key mechanotransducers of hemodynamic forces(*4*). Under LSS, ECs adopt an atheroprotective phenotype marked by increased nitric oxide production, anti-inflammatory signaling, and preserved barrier integrity. In contrast, OSS drives pro-atherogenic responses, including enhanced monocyte recruitment, oxidative stress, and compromised barrier function, thereby initiating early atherosclerotic plaque formation(*5–7*). Unraveling the key endothelial sensors that respond to variations in blood flow shear stress is crucial for the development of targeted therapies aimed at preventing lesion formation at predisposed arterial sites (*8*).

A key mechanism underlying ECs mechanotransduction of shear stress involves post-translational modifications (PTMs) of proteins, which enable rapid and reversible cellular adaptations(*9–13*). Neddylation is a reversible post-translational modification similar to ubiquitination. In this process, the ubiquitin-like protein NEDD8 is attached to lysine residues on target proteins, mainly cullins family proteins. This process is mediated by a three-step enzymatic cascade involving a single E1-activating enzyme (the NAE1 [NEDD8-activating enzyme E1 subunit 1]/NAE2 [NEDD8-activating enzyme E1 subunit 2] complex), two E2-conjugating enzymes (UBE2M [ubiquitin conjugating enzyme E2 M] and UBE2F [ubiquitin conjugating enzyme E2 F]), and multiple substrate-specific E3 ligases(*14, 15*). Neddylation regulates the activity, stability, and function of modified proteins(*16, 17*). Inflammatory stimuli upregulate endothelial neddylation(*18*), and mechanical cues, such as cyclic stretch, have similarly been shown to increase neddylation levels in pulmonary endothelial cells(*19*). However, the functional role of neddylation in endothelial responses to shear stress, particularly in the context of disturbed flow-induced atherogenesis, remains largely unexplored.

The mortality factor 4-like protein 1 (MORF4L1, also called MRG15) is a highly conserved chromatin-binding protein. MRG15 serves as a core scaffold subunit in multiple histone acetylation complexes, including the NuA4/TIP60 acetyltransferase complex and Sin3-HDAC deacetylase complex(*20–22*). It plays a vital role in epigenetic regulation, DNA damage repair, cell proliferation, senescence, and apoptosis(*23–26*). MRG15 is indispensable for normal embryonic development, as its global knockout results in sublethal embryonic phenotypes in mice, with only rare instances of perinatal survival. Histological examination of *Mrg15*^−/−^ embryos at embryonic day (E) 14 reveals severe vascular abnormalities, including disrupted vascular patterning and impaired vessel maturation(*27*). Genetic knockout studies targeting cardiac progenitor cells (CPCs) and cardiomyocytes demonstrate that MRG15 regulates cardiomyocyte proliferation during heart development and following injury(*28*). Large-scale genome-wide association studies have identified multiple coronary artery disease risk loci at or near the MRG15 locus(*29–31*), yet its mechanistic contribution to atherosclerosis is incompletely understood.

In the present study, we investigated the role of endothelial MRG15 in disturbed flow-induced atherosclerosis using partial carotid ligation (PCL) and chronic Western diet-fed mouse models, aiming to elucidate its function in OSS-mediated endothelial activation and identify potential therapeutic strategies for atherosclerosis.

## Material and methods

### Human artery sample

All procedures involving human samples were conducted in accordance with the ethical principles outlined in the Declaration of Helsinki. Human arterial samples were obtained from patients with or without atherosclerosis at Zhongshan Hospital, Fudan University. Written informed consent was obtained from all participants. Aortic tissue samples were fixed in formalin, paraffin-embedded, and serially sectioned for subsequent histological examination. This study was approved by the Ethics Committee of Zhongshan Hospital.

### Animal Studies

All animal experiments were approved by the Institutional Animal Care and Use Committee (IACUC) of the School of Basic Medical Sciences, Fudan University, and were performed in strict compliance with the National Institutes of Health (NIH) Guide for the Care and Use of Laboratory Animals.

EC-specific *Mrg15*-inducible knockout mice (*Mrg15*^iECKO^) were generated by crossing transgenic mice expressing Cre-recombinase specifically in EC (*Cdh5-Cre*^ERT2^) with *Mrg15*^fl/fl^ mice. *Mrg15*^fl/fl^ mice were generated by flanking exons2 of *Mrg15* with loxP sites from Shanghai Model Organisms Center, Inc. 6-week-old male *Mrg15*^fl/fl^ mice and *Mrg15*^iECKO^ littermates (20-25 g; n=6) were intraperitoneally injected with 100 mg/kg of tamoxifen resolved in corn oil every other day for a total of four times.

EC-specific *Mrg15*-inducible overexpression mice (*Mrg15*^iECTG^) were generated by crossing transgenic mice expressing Cre-recombinase specifically in EC (*Cdh5-Cre*^ERT2^) with *Mrg15*^WT^ mice. Using Clustered Regularly Interspaced Short Palindromic Repeats (CRISPR)/Cas9-mediated homologous recombination, the loxP-STOP-loxP-Mrg15-2A-tdTomato expression cassette was inserted into the *Rosa26* locus to create the *Rosa26-Mrg15*^TG^(*Mrg15*^WT^) mice from Shanghai Model Organisms Center, Inc. 6-week-old male *Mrg15*^WT^ mice and *Mrg15*^iECTG^ littermates (20-25 g; n=6) were intraperitoneally injected with 100 mg/kg of tamoxifen resolved in corn oil was injected intraperitoneally for four every other day to CDH5 every other day for a total of four times.

### Partial carotid ligation model

The partial carotid ligation (PCL) model is a well-established approach to induce disturbed blood flow and accelerate atherosclerosis in vivo(*32*). To establish hypercholesterolemia, mice received tail-vein injection of AAV8-PCSK9 gain-of-function virus (1 × 10^11^ viral genomes) to induce liver-specific low-density lipoprotein receptor (LDLR) knockdown, followed by feeding with Western diet (Shuyishuer Biotechnology Co. Ltd, Changzhou, China, D12109C). For surgery, mice were anesthetized with an intraperitoneal injection of ketamine (100 mg/kg) and xylazine (10 mg/kg). A 4–5 mm midline vertical incision was made in the neck, and the left common carotid artery and its branches were exposed by blunt dissection. The left external carotid, internal carotid, and occipital arteries were permanently ligated with 6-0 silk sutures, while the superior thyroid artery was left patent to preserve residual flow. Postoperatively, animals were placed on a heating pad and monitored until complete recovery from anesthesia. All surgical procedures were performed by the same investigator to minimize inter-operator variability.

### Shear stress study in vitro

In vitro shear stress experiments were conducted as previously described(*10*). Briefly, human umbilical vein endothelial cells (HUVECs; 5 × 10^4^ cells) were seeded onto slides. The slides were assembled into flow chambers and connected to the Ibidi flow system (IBIDI, Germany). Cells were then subjected to LSS at 12 dyn/cm^2^ or OSS at 6 ± 0.5 dyn/cm^2^ with a 1 Hz frequency. The entire flow system was maintained within a humidified incubator at 37°C with 5% CO₂.

### RNA Extraction and RT-qPCR for gene expression

Total RNA was extracted from HUVECs, human carotid plaques, primary mouse endothelial cells, or mouse tissues using TRIzol Reagent (Invitrogen, Carlsbad, CA, USA; cat. no. 15596026). Reverse transcription was performed using the High-Capacity cDNA Reverse Transcription Kit (Toyobo). Quantitative real-time PCR (RT-qPCR) was conducted on a Bio-Rad iQ5 thermal cycler with SYBR Green Supermix (Toyobo, Osaka, Japan; cat. no. QPK-201). The amplification protocol consisted of an initial denaturation step at 95°C for 5 min, followed by 40 cycles of 95°C for 15 s, 58°C for 30 s, and 72°C for 30 s. A melting curve analysis was performed from 60°C to 95°C with a ramp rate of 1°C/min to confirm specificity. Relative gene expression was calculated using the 2^−ΔΔCt^ method and normalized to the housekeeping gene β-actin.

### Immunoblotting

Cells were lysed in ice-cold buffer containing 20 mmol/L Tris-HCl (pH 7.5), 150 mmol/L NaCl, 1% Triton X-100, 1 mmol/L EDTA, 1 mmol/L EGTA, 2.5 mmol/L sodium pyrophosphate, 1 mmol/L β-glycerophosphate, 50 mmol/L NaF, 1 mmol/L Na₃VO₄, and protease inhibitor cocktail. Following centrifugation at 12,000 rpm for 15 min at 4°C, the supernatant was collected for western blot analysis as previously described. Equal amounts of protein were separated by SDS-PAGE and transferred to PVDF membranes. Membranes were blocked and then incubated overnight at 4°C with primary antibodies against MRG15 (Cell Signaling Technology, #14098), ITGA5 (Proteintech, #10569-1-AP), ICAM1 (Cell Signaling Technology, #4915), β-actin (Proteintech, #66069-1-Ig), and GAPDH (Proteintech, #60004-1-Ig). After three washes with 1× TBST, membranes were incubatedwith appropriate HRP-conjugated secondary antibodies for 2 h at room temperature. Immunoreactive bands were visualized using an Odyssey Infrared Imaging System (LI-COR Biosciences). Densitometric quantification was performed using NIH ImageJ software, with total protein levels normalized to β-actin.

### En face staining and immunofluorescence

Mice were euthanized by CO₂ inhalation, after which the thoracic aorta and carotid arteries were carefully dissected. Perivascular connective tissue and fat were removed under a stereomicroscope. For immunofluorescence, cryosections, cultured cells, and intact arterial segments were fixed in 4% paraformaldehyde for 15 min at room temperature and permeabilized with 0.1% Triton X-100 in PBS for 10 min. Samples were blocked with 5% bovine serum albumin (BSA) in PBS for 1 h at room temperature, then incubated overnight at 4°C (or 48 h for intact arteries) with primary antibodies diluted in blocking buffer. Following three washes with PBS, specimens were incubated with fluorochrome-conjugated secondary antibodies for 1 h at room temperature in the dark. Samples were mounted with anti-fade mounting medium (Vector Laboratories) and imaged using a TCS SP8 inverted confocal microscope (Leica Microsystems, Germany).The antibodies used in this study include anti-MRG15 (1:100, HPA062010, Sigma); anti- CD31 (1:100, #AF3628, R&D systems); anti-ICAM1 (1:200, #ab282575, Abcam); anti-CD68 (1:250, #A24386PM, Abclonal); Donkey anti-Goat IgG (H+L) Highly Cross-Adsorbed Secondary Antibody, Alexa Fluor™ Plus 647 (1:1000, # A32849, Invitrogen); Donkey anti-Rabbit IgG (H+L) Highly Cross-Adsorbed Secondary Antibody, Alexa Fluor™ Plus 488 (1:1000, #A32790, Invitrogen), Donkey anti-Rabbit IgG (H+L) Highly Cross-Adsorbed Secondary Antibody, Alexa Fluor™ Plus 594 (1:1000, #A32754, Invitrogen). To validate antibody specificity and distinguish genuine target staining from the background, secondary antibody-only was used as a control. Representative images were chosen based on group mean/average and image quality.

### Immunoprecipitation

Whole-cell lysates were prepared for immunoprecipitation. Cells were lysed in ice-cold buffer containing 20 mmol/L Tris-HCl (pH 7.5), 150 mmol/L NaCl, 1% Triton X-100, 1 mmol/L EDTA, 1 mmol/L EGTA, 2.5 mmol/L sodium pyrophosphate, 1 mmol/L β-glycerophosphate, 50 mmol/L NaF, 1 mmol/L Na₃VO₄, and protease inhibitor cocktail. Lysates were incubated overnight at 4°C with rotation using the mouse monoclonal anti-MRG15 antibody. The following day, immune complexes were captured by adding Protein A/G PLUS-Agarose (Santa Cruz Biotechnology, Dallas, TX, USA; cat. no. sc-2003) and incubating for 3 h at 4°C. Beads were washed three times with lysis buffer, and bound proteins were eluted by boiling in 1× SDS sample buffer at 100°C for 10 min. Immunoblotting was subsequently performed following standard procedures.

### Adhesion Assay

Monocyte adhesion to endothelial cells (ECs) was assessed as previously described (*33*). Briefly, ECs were seeded in 96-well plates and cultured to 70–80% confluence. THP-1 monocytes were labeled with PKH26 red fluorescent dye using the PKH26 Red Fluorescent Cell Linker Mini Kit according to the manufacturer’s protocol. Labeled THP-1 cells were then added to the EC monolayers and incubated for 30 min at 37°C. Non-adherent cells were removed by gentle washing with PBS, and the number of adherent fluorescent cells was quantified in randomly selected fields under a fluorescence microscope.

### Analysis of scRNA-sequencing data

The Seurat (version 5.0) R package(*34*) was used to quality filter and cluster cells. Then, doublets were identified using DoubletFinder(*35*) and removed from downstream analysis. Pseudotime analysis was performed by the Monocle 3 package(*36*), as described in the github (https://cole-trapnell-lab.github.io/monocle3/). EC were isolated and used as the input expression matrix for trajectory analysis. Transcription factor activities were calculated by DoRothEA (https://github.com/saezlab/dorothea)(37). Gene Ontology analysis(*38*) and KEGG Pathway analysis(*39*) were performed on David(https://david.ncifcrf.gov/)(40).

### Chromatin immunoprecipitation (ChIP)

ChIP were performed as previously described(*41*). Briefly, cells were cross-linked with 1% formaldehyde for 10 min at room temperature and quenched with 0.125 M glycine for 10 min. Cell pellets were resuspended in low-salt lysis buffer (50 mM HEPES/KOH pH 7.5, 150 mM NaCl, 1% Triton X-100, 0.05% SDS, 10 mM EDTA, supplemented with protease inhibitors) and sonicated using a Q800R2 DNA shearing sonicator (Qsonica LLC, Newtown, CT) for 20 min (30-s on/off cycles) to generate chromatin fragments of 200–400 bp. Immunoprecipitation was performed by incubating the sonicated chromatin overnight at 4°C with anti-EZH2 or anti-H3K27me3 antibodies, followed by capture with Protein A/G PLUS-Agarose beads. Beads were sequentially washed with low-salt lysis buffer and low-salt wash buffer (10 mM Tris-HCl, pH 8.0, 250 mM LiCl, 1 mM EDTA, 0.5% NP-40, 0.5% sodium deoxycholate). Bound chromatin was eluted in elution buffer (50 mM Tris-HCl pH 8.0, 10 mM EDTA, 1% SDS) and treated with Proteinase K (Yeasen, Shanghai, China; cat. no. 10401ES60) and RNase A (Thermo Fisher, Waltham, MA; cat. no. EN0531) at 65°C for 4 h. Cross-linking was reversed, and DNA was purified for downstream analysis.

### Assay for transposase-accessible chromatin using sequencing (ATAC-seq)

Assay for transposase-accessible chromatin using sequencing (ATAC-seq) was performed as previously described(*42*). Briefly, cells were detached with trypsin (Invitrogen, Carlsbad, CA; cat. no. 25200072) and pretreated with 200 U/mL DNase I (Sigma-Aldrich, St. Louis, MO; cat. no. 260913) for 30 min at 37°C to eliminate extracellular and dead-cell DNA. A total of 50,000 viable cells were resuspended in 1 mL cold RSB buffer (10 mM Tris-HCl, pH 7.4, 10 mM NaCl, 3 mM MgCl₂) and centrifuged at 500 × g for 5 min. Nuclei were isolated by resuspending the pellet in 50 μl RSB buffer supplemented with 0.5% NP-40, 0.1% Tween-20, and 0.01% digitonin, followed by incubation on ice for 10 min. After lysis, 1 mL of RSB buffer containing 0.1% Tween-20 was added, and nuclei were pelleted by centrifugation at 500 × g for 10 min. ATAC-seq libraries were then prepared using the TruePrep DNA Library Prep Kit V2 for Illumina (Vazyme Biotech, Nanjing, China; cat. no. TD501-01) according to the manufacturer’s instructions. Sequencing was performed on a NovaSeq 6000 platform by Annoroad Gene Technology.

### Statistical Analysis

Data are expressed as mean ± standard error of the mean (SEM). Normality of data distribution was evaluated using the Shapiro–Wilk test. For comparisons between two groups with normally distributed data, we applied the two-tailed unpaired Student’s t-test (assuming equal variances) or Welch’s t-test (when unequal variances were confirmed by an F-test). For comparisons involving three or more groups, variance homogeneity was first assessed with the Brown–Forsythe test. When variances were similar, one-way ANOVA followed by Tukey’s multiple comparisons test was performed. Non-normally distributed data were analyzed using the Mann–Whitney U test for two-group comparisons or the Kruskal–Wallis test followed by Dunn’s post hoc test for multiple groups. Detailed statistical analyses are provided in the corresponding figure legends. The number of biological replicates for each group is indicated in the figure legends. Representative images were chosen to best represent the group mean from all available data. Statistical significance was defined as p < 0.05. All statistical analyses were conducted using GraphPad Prism version 8.0 (GraphPad Software, San Diego, CA, USA).

## Results

### Atherogenic Shear Stress Suppresses Endothelial MRG15 Expression

To identify potential therapeutic targets for atherosclerosis, we performed an integrative analysis of downregulated proteins in proteomic profiles from human umbilical vein endothelial cells (HUVECs) under disturbed flow and coronary artery disease risk-associated genes derived from large-scale genome-wide association studies (GWAS)(*43*). Intersection analysis of these datasets yielded nine candidate proteins, of which only two proteins displayed significant downregulation: TIMP2 (tissue inhibitor of metalloproteinase 2) and MRG15 (Figure 1A). TIMP2 has been established as a pro-atherogenic gene (*44*). However, whether MRG15 responds to flow shear stress stimulation and its role in atherosclerosis remain unclear. To investigate the hemodynamic modulation of endothelial MRG15, we compared its protein levels by western blot between two distinct mouse aortic regions: the aortic arch (AA) and the thoracic aorta (TA). The results demonstrated that MRG15 levels were significantly lower in the AA, which is constantly exposed to disturbed flow, compared with the TA exposed to unidirectional laminar flow (Figure 1B). En face immunofluorescence staining of the mouse aortic endothelium showed consistent results that the inner curvature of the aortic arch expressed a lower level of MRG15 than the thoracic aorta (Figure 1C). To further verify flow-dependent regulation of MRG15 expression in vivo, we induced disturbed flow in the left carotid artery (LCA) of mice via PCL for 7 days (Figure 1D). En face immunofluorescence staining revealed decreased MRG15 protein expression in the aortic endothelium of the LCA following PCL (Figure 1D). To determine the expression of MRG15 in ECs in response to different types of shear stress generated by blood flow, HUVECs were exposed for different durations to OSS or LSS generated by the Ibidi pump system. Gene expression results showed that, compared with cells kept in STA, MRG15 protein expression was inhibited by OSS (Figure 1E and Figure 1G). By contrast, LSS upregulated MRG15 protein (Figure 1F). Notably, MRG15 was downregulated in the endothelium of lesion areas in human atherosclerotic arteries (Figure 1H). In the mouse carotid artery plaque, endothelial expression of MRG15 was found to be decreased (Figure 1I). Collectively, these results identify MRG15 as a flow-regulated gene that may contribute to the maintenance of endothelial homeostasis and atheroprotection.

**Fig. 1.**
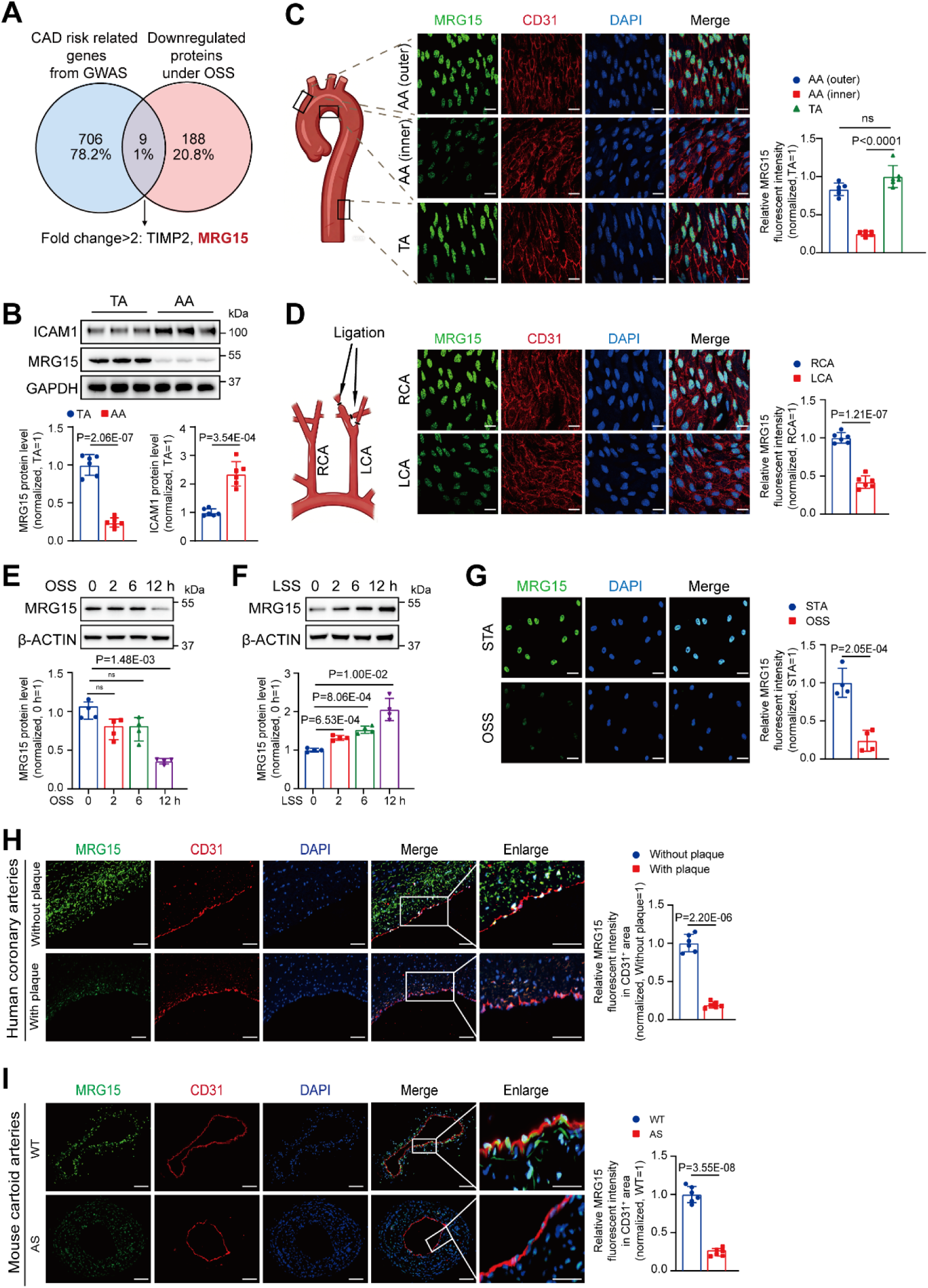
**Disturbed flow downregulates the expression of MRG15 in the endothelium.** (A) Venn diagram of downregulated proteins from comparative proteomic profiling of human umbilical vein endothelial cells (HUVECs) under oscillatory shear stress (OSS) 12 hours (n=4) compared to static (STA) (n=4), and genes associated with or located near single-nucleotide polymorphisms (SNPs) linked to increased risk of coronary artery disease (CAD). The selection criteria for the proteomic data set were fold change >1.5 and adj.P.Val<0.05. (B) Western blot analysis of MRG15, ICAM1, and GAPDH levels in the aortic arch (AA) and the thoracic aorta (TA) of the mice (n=6). (C) En-face immunofluorescence staining for MRG15 (green), CD31 (red), and 4′,6-diamidino-2-phenylindole (DAPI, blue) in the inner or outer of the (AA) and the TA of the mice (n=6) (Scale bar, 50 μm). (D) En-face immunofluorescence staining for MRG15 (green), CD31 (red), and DAPI (blue) in the carotid arteries of mice at 4 weeks after partial carotid ligation (PCL) surgery (n=6) (Scale bar, 50 μm). (E) Western blot analysis of MRG15, ICAM1, and β-ACTIN levels in HUVECs subjected to STA or OSS (6 ± 0.5 dyn/cm2; 1 Hz; n=4) at the indicated times. (F) Western blot analysis of MRG15, ICAM1, and β-ACTIN levels in HUVECs subjected to STA or laminar shear stress (LSS) (12 dyn/cm^2^; n=4) at the indicated times. (G) Immunofluorescence staining for MRG15 (green), DAPI (blue) in HUVECs subjected to STA or OSS (6 ± 0.5 dyn/cm^2^; 1 Hz; n=4) (Scale bar, 20 μm). (H) Immunofluorescence staining for MRG15 (green), CD31 (red), and DAPI (blue) in the human coronary arteries atherosclerotic lesions (n=6) and healthy human coronary arteries (n=6) (Scale bar, 20 μm). (I) Immunofluorescence staining for MRG15 (green), CD31 (red), and DAPI (blue) in the mice carotid aortic atherosclerotic lesions (n=6) and normal carotid aortas (n=6). The statistical analysis was performed by two-tailed Student t test for D, G, I, and MRG15 in B, by Welch’s t test for ICAM1 in B and H, by Brown-Forsythe tests for C, E, and F.

### Disturbed Flow Accelerates MRG15 Degradation by Enhancing Neddylation

Next, we found that OSS stimulation did not alter *MRG15* mRNA levels (Figure 2A), indicating that the downregulation is likely mediated by post-transcriptional mechanisms. To assess the impact of OSS on MRG15 stability, we conducted cycloheximide (CHX) chase experiments and observed a markedly shortened half-life of MRG15 under OSS 6 hours relative to STA (Figure 2B). This result indicates that OSS accelerates MRG15 degradation, thereby decreasing its protein stability. Furthermore, OSS stimulation markedly increased the overall neddylation levels in endothelial cells (Figure 2C). Concurrently, we found that OSS stimulation significantly downregulated MRG15 expression in HUVECs. This effect was effectively reversed by the neddylation inhibitor MLN4924 (Figure 2D), suggesting that disturbed flow promotes neddylation as a mechanism contributing to MRG15 destabilization.

**Fig. 2.**
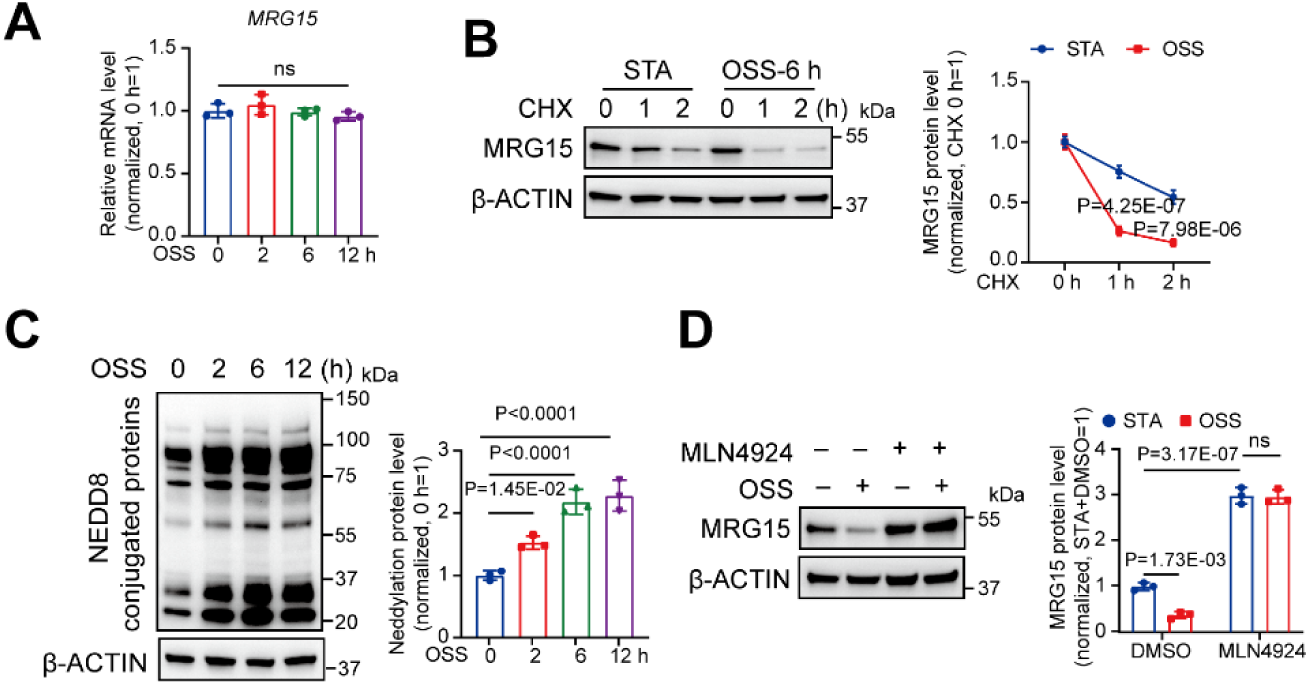
**Disturbed Flow Accelerates MRG15 Degradation by Enhancing Neddylation.** (A) Real time-qPCR analysis of the expression of MRG15 in human umbilical vein endothelial cells (HUVECs) subjected to oscillatory shear stress (OSS) (6 ± 0.5 dyn/cm^2^; 1 Hz; n=4) at the indicated times (n=3). (B) Western blot analysis of protein levels of MRG15 and β-ACTIN. HUVECs were cultured under STA or OSS for 6 hours and treated with DMSO or CHX (10 μg/ml) at the indicated times before harvesting (n=3). (C) Western blot analysis of protein levels of global NEDD8 conjugation and β-ACTIN in HUVECs subjected to STA or OSS (6 ± 0.5 dyn/cm^2^; 1 Hz; n=3) at the indicated times. (D) Western blot analysis of protein levels of MRG15 and β-ACTIN. HUVECs were cultured under STA or OSS for 12 hours and treated with DMSO or MLN4924 (1 μg/ml) for 8 hours (n=3). The statistical analysis was performed by Ordinary one-way ANOVA test for A and C, by two-way ANOVA followed by Tukey’s post hoc test for B and D.

### EC-Specific *Mrg15* Deletion Aggravates Atherosclerosis in Mice

Next, we investigated the role of endothelial Mrg15 in atherosclerosis using the PCL mouse model, a well-established approach for studying flow-dependent atherogenesis. We employed the Cre-loxP system to generate mice with inducible, endothelial-specific deletion of *Mrg15* (*Mrg15*^iECKO^). This was achieved by crossing *Mrg15*^fl/fl^ mice, where loxP sites flank exon 2, with *Cdh5-CreERT2* mice that express tamoxifen-inducible Cre recombinase in endothelial cells. The knockout efficiency of MRG15 was verified by immunofluorescent staining in the vascular endothelium (Figure 3A). Seven-week-old male *Mrg15*^fl/fl^ mice and *Mrg15*^iECKO^ mice were infected with AAV8-*Pcsk9*^D377Y^ and subsequently subjected to partial left carotid artery ligation following four weeks of Western diet (Figure 3B). We found that EC-specific *Mrg15* knockout led to an increase in the formation of atherosclerotic plaques induced by disturbed flow (Figure 3C and Figure 3D). Immunofluorescence staining showed an enhanced infiltration of CD68^+^ macrophages. (Figure 3E). The pro-atherogenic role of EC-specific knockout of *Mrg15* was further validated in a Western diet-induced hyperlipidemia model. We infected both *Mrg15*^flox/flox^ and *Mrg15*^iECKO^ mice with AAV8-*Pcsk9*^D377Y^. Twelve weeks of the Western diet was administered to induce hypercholesterolemia (Figure 3F). En face analysis of aortae from mice revealed that EC-specific knockout of *Mrg15* exacerbated plaque deposition (Oil Red O; Figure 3G). Consistently, cross-sectional assessment of aortic roots showed lipid accumulation (Oil Red O; Figure 3H) and increased the infiltration of CD68^+^ macrophages within the lesions (Figure 3I), confirming the acceleration of atherosclerosis. Collectively, these data demonstrate that depletion of endothelial *Mrg15* aggtavates atherosclerotic lesions in mice.

**Fig. 3.**
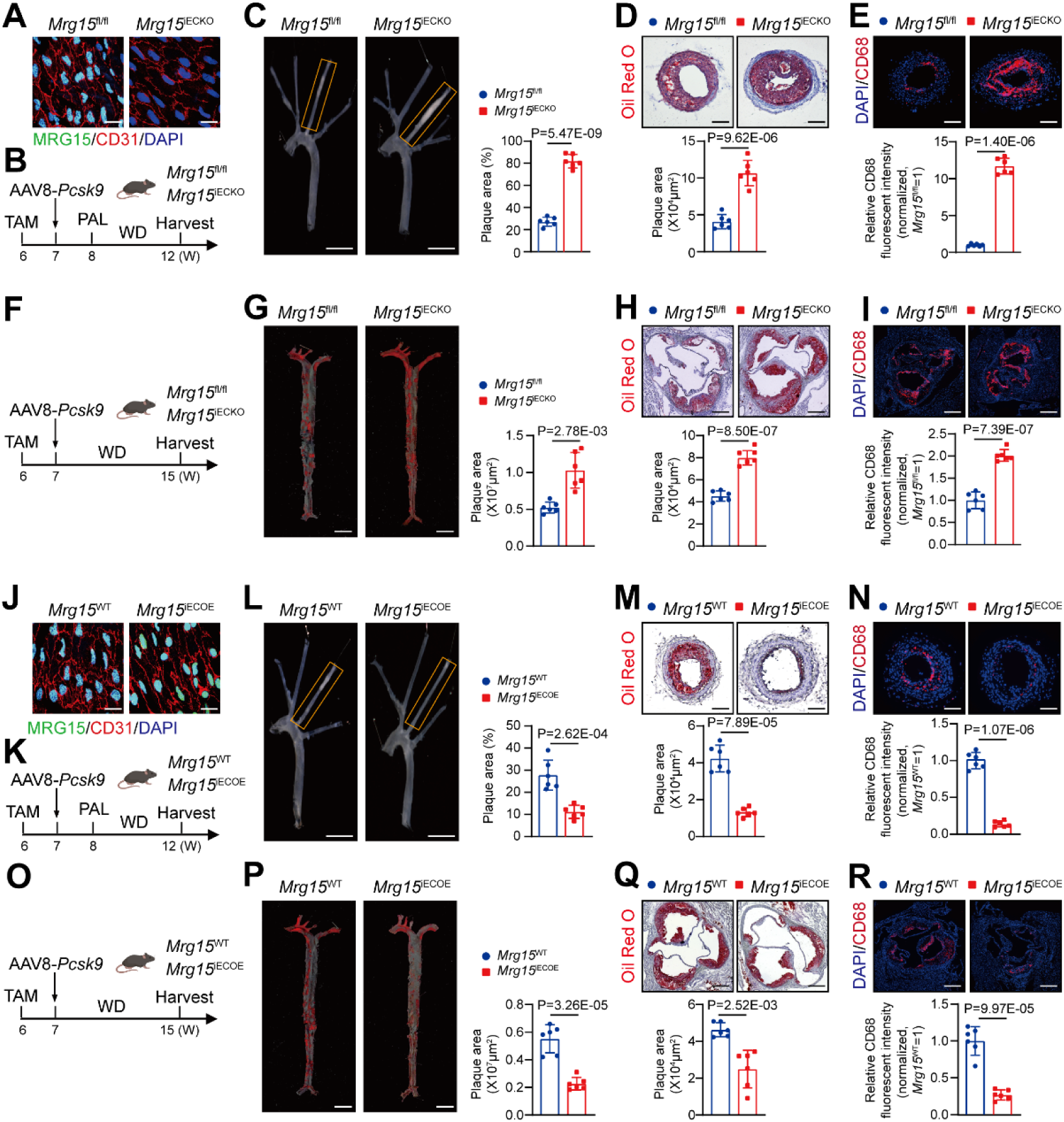
**Endothelial *Mrg15* Deletion Exacerbates Atherosclerosis, While Overexpression Attenuates It** (A) En face immunofluorescence staining for MRG15 (green), CD31 (red), and 4′,6-diamidino-2-phenylindole (DAPI, blue) in the arterial endothelium of *Mrg15*^fl/fl^ and *Mrg15*^iECKO^ mice (Scale bar, 50 µm). (B) The study design employing a partial carotid ligation model. (C) Representative images and quantitative analysis of plaque areas of ligated carotid arteries from the mice in *Mrg15*^fl/fl^ mice (n=6) and *Mrg15*^iECKO^ mice (n=6) (Scale bar, 1 mm). (D) Representative result and quantitative analysis of Oil Red O staining of ligated left carotid artery (LCA) sections from the mice in *Mrg15*^fl/fl^ mice (n=6) and *Mrg15*^iECKO^ mice (n=6) (Scale bar, 50 µm). (E) Representative result and quantitative analysis of CD68 staining of LCA sections from the mice in *Mrg15*^fl/fl^ mice (n=6) and *Mrg15*^iECKO^ mice (n=6) (Scale bar, 50 µm). (F) The study design using the chronic high cholesterol diet-induced atherosclerotic model. (G) Representative result and quantitative analysis of Oil Red O staining of the aortae from the mice in *Mrg15*^fl/fl^ mice (n=6) and *Mrg15*^iECKO^ mice (n=6) (Scale bar, 1 mm). (H) Representative image and quantitative analysis of aortic sinus lesion with Oil Red O staining in aortic roots from the mice in *Mrg15*^fl/fl^ mice (n=6) and *Mrg15*^iECKO^ mice (n=6) (Scale bar, 50 µm). (I) Representative image and quantitative analysis of aortic sinus lesion with CD68 staining in aortic roots from the mice in *Mrg15*^fl/fl^ mice (n=6) and *Mrg15*^iECKO^ mice(n=6) (Scale bar, 50 µm). (J) En face immunofluorescence staining for MRG15 (green), CD31 (red), and DAPI (blue) in the arterial endothelium of *Mrg15*^WT^ and *Mrg15*^iECOE^ mice (Scale bar, 50 µm). (K) The study design employing a partial carotid ligation model. (L)Representative images and quantitative analysis of plaque areas of ligated carotid arteries from the mice in *Mrg15*^WT^ mice (n=6) and *Mrg15*^iECOE^ mice (n=6) (Scale bar, 1 mm). (M) Representative result and quantitative analysis of Oil Red O staining of LCA sections from the mice in *Mrg15*^WT^ mice (n=6) and *Mrg15*^iECOE^ (n=6) (Scale bar, 50 µm). (N) Representative result and quantitative analysis of CD68 staining of LCA sections from the mice in *Mrg15*^WT^ mice (n=6) and *Mrg15*^iECOE^ mice(n=6) (Scale bar, 50 µm). (O) The study design using the chronic high cholesterol diet-induced atherosclerotic model. (P) Representative result and quantitative analysis of Oil Red O staining of the aortae from the mice in *Mrg15*^WT^ mice (n=6) and *Mrg15*^iECOE^ mice (n=6) (Scale bar, 1 mm). (Q) Representative image and quantitative analysis of aortic sinus lesion with Oil Red O staining in aortic roots from the mice in *Mrg15*^WT^ mice (n=6) and *Mrg15*^iECOE^ mice (n=6) (Scale bar, 50 µm). (R) Representative image and quantitative analysis of aortic sinus lesion with CD68 staining in aortic roots from the mice in *Mrg15*^WT^ mice (n=6) and *Mrg15*^iECOE^ mice(n=6) (Scale bar, 50 µm). The statistical analysis was performed by two-tailed Student t test for C, D, H, I, L, P , by Welch’s t test for E, G, M, N, P, Q, R.

### EC-Specific *Mrg15* Overexpression Attenuates Atherosclerosis in Mice

To determine whether endothelial MRG15 exerts an anti-atherosclerotic effect with translational therapeutic potential, we generated endothelial-specific *Mrg15*-overexpressing mice using the Cre-loxP system. Using CRISPR/Cas9-mediated homologous recombination, the loxP-STOP-loxP-*Mrg15*-2A-tdTomato expression cassette was inserted into the *Rosa26* locus to create the *Rosa26*-*Mrg15*^TG^ mice. The *Rosa26*-*Mrg15*^TG^ mice were bred with *Cdh5-CreERT2* mice to create mice with overexpression of *Mrg15* in ECs (*Mrg15*^iECOE^) upon tamoxifen induction. The overexpression efficiency of MRG15 was verified by immunofluorescent staining in the vascular endothelium (Figure 3J). Seven-week-old male *Mrg15*^WT^ and *Mrg15*^iECOE^ mice were infected with AAV8-*Pcsk9*^D377Y^ and subsequently subjected to partial left carotid artery ligation following four weeks of Western diet (Figure 3K). Our results showed that endothelium-specific overexpression of *Mrg15* decreased disturbed flow-induced atherosclerosis (Figure 3L and Figure 3M). Immunofluorescence analysis of CD68 in ligated carotid artery sections revealed significantly attenuated infiltration of CD68^+^ macrophages in mice with endothelial-specific *Mrg15* overexpression (Figure 3N). We found that endothelial-specific *Mrg15* overexpression significantly reduced atherosclerotic plaque formation in mice fed the high-cholesterol diet, as assessed by en face Oil Red O staining of the aorta (Figure 3O and Figure 3P). This protective effect was further confirmed by Oil Red O staining of aortic root sections (Figure 3Q). CD68 immunofluorescence staining of the aortic roots also demonstrated that endothelium - specific overexpression of *Mrg15* reduced macrophage infiltration in atherosclerotic plaques (Figure 3R). Taken together, our collective in vivo evidence from both loss-of-function and gain-of-function studies establishes a suppressive role for MRG15 in atherogenesis.

### *Mrg15* deficiency primarily affects endothelial cell dynamics in mouse atherosclerosis

To explore the mechanism of endothelial MRG15 suppressing atherosclerosis in an unbiased manner, we performed scRNA-seq analysis of LCA from *Mrg15*^flox/flox^ and *Mrg15*^iECKO^ mice (n=6) by PCL following two weeks of western diet. A total of 6614 and 8513 cells were captured in LCA from *Mrg15*^fl/fl^ and *Mrg15*^iECKO^ mice, respectively. Unsupervised clustering of the scRNA-seq identified seven cell clusters, including smooth muscle cell (SMC), macrophage, EC, fibroblast, adipocyte, neutrophil, and schwann cell (Figure 4A). A marked increase in the macrophage proportion was observed in the atherosclerotic carotids of *Mrg15*^iECKO^ mice compared to those from *Mrg15*^fl/fl^ mice (Figure 4B). Transcription factor (TF) activity profiling in the EC cluster revealed significant activation of pro-atherogenic transcription factors in *Mrg15*^iECKO^ mice, including *Jun*, *Snail2*, and *Bach1*, alongside concomitant suppression of atheroprotective transcription factors, such as *Erg*, *Klf4*, and *Epas1* (Figure 4C). Gene ontology (GO) enrichment analysis revealed significant activation of biological processes related to positive regulation of immune effector process and cell adhesion mediated by integrin in the *Mrg15*^iECKO^ group (Figure 4D). Similarly, Kyoto Encyclopedia of Genes and Genomes (KEGG) pathway analysis demonstrated enrichment of pathways associated with fluid shear stress and atherosclerosis, as well as the TNF signaling pathway, in the *Mrg15*^iECKO^ group (Figure 4E).

**Fig. 4.**
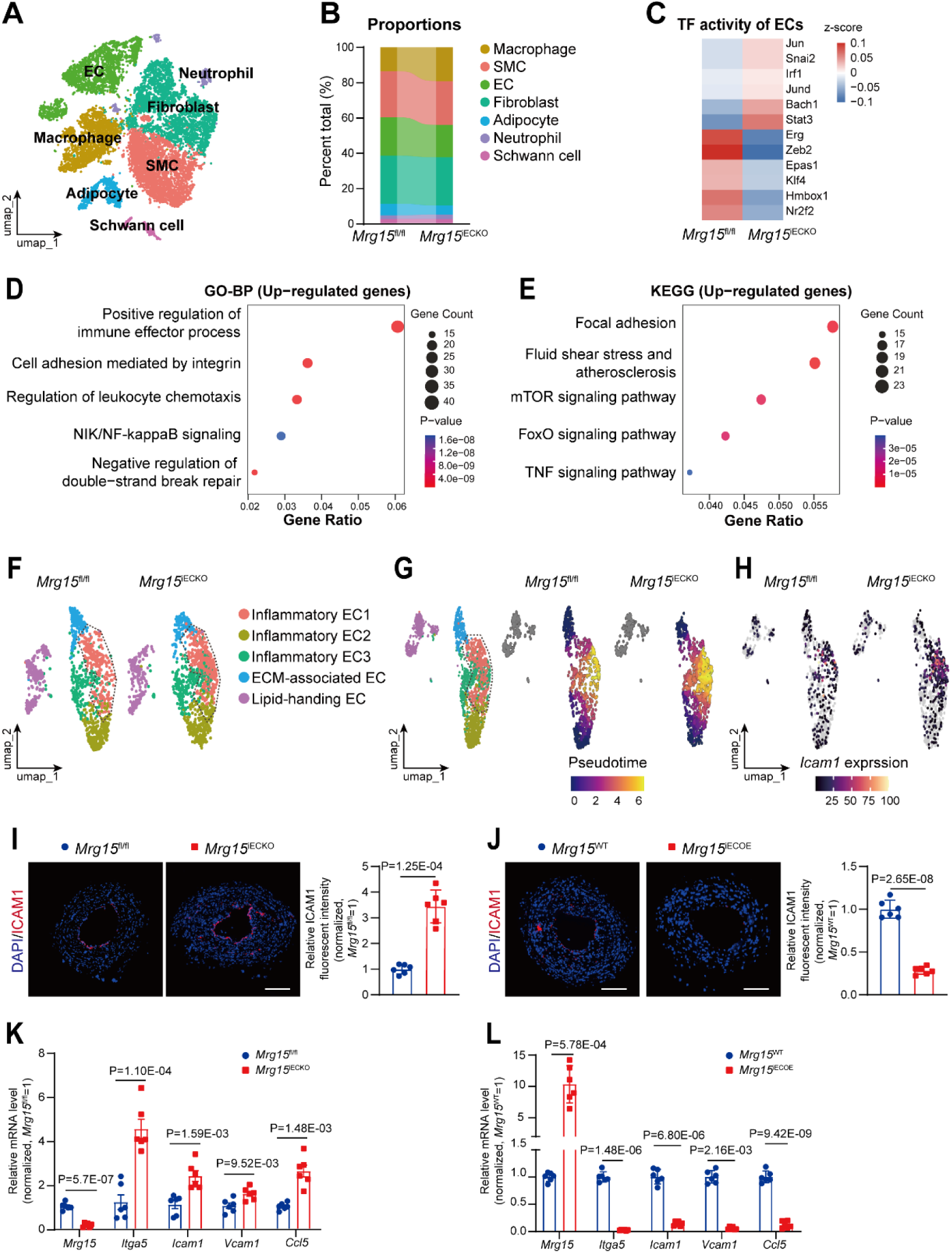
**Endothelial *Mrg15* Deletion promotes inflammation in atherosclerosis.** (A) UMAP plots showed Single-cell RNA sequencing (scRNA-seq) analyzed in the carotid arteries of mice at 4 weeks after partial carotid ligation (PCL) surgery from the mice in *Mrg15*^fl/fl^ mice (n=6) and *Mrg15*^iECKO^ mice(n=6), color-coded for cell types. (B) The proportion of cell type in *Mrg15*^fl/fl^ mice and *Mrg15*^iECKO^ mice. (C) Heatmap of selected tran transcription factor (TF) activities inferred with DoRothEA from gene expression data generated in EC from *Mrg15*^fl/fl^ mice and *Mrg15*^iECKO^ mice. (D) Gene Ontology analysis of upregulated genes in EC from *Mrg15*^iECKO^ mice compared to *Mrg15*^fl/fl^ mice. (E) Kyoto Encyclopedia of Genes and Genomes analysis of upregulated genes in EC from *Mrg15*^iECKO^ mice compared to *Mrg15*^fl/fl^ mice. (F) UMAP plots showed single-cell transcriptomes analyzed in EC from *Mrg15*^fl/fl^ mice and *Mrg15*^iECKO^ mice, color-coded for cell types, split by sample origins. (G) Left: Trajectory analysis in EC UMAPs demonstrated EC subtype. Middle and right: Trajectory analysis in EC UMAPs demonstrates pseudo-time, split by sample origins. (H) Trajectory analysis in EC UMAPs demonstrated *Icam1* expression, split by sample origins. (I) Representative result and quantitative analysis of ICAM1 staining of LCA sections from the mice in *Mrg15*^fl/fl^ mice (n=6) and *Mrg15*^iECKO^ mice(n=6) (Scale bar, 50 µm). (J) Representative result and quantitative analysis of ICAM1 staining of LCA sections from the mice in *Mrg15*^WT^ mice (n=6) and *Mrg15*^iECOE^ mice(n=6) (Scale bar, 50 µm). (K) Real time-qPCR analysis of the expression of *Mrg15*, *Itga5*, *Icam1*, *Vcam1* and *Ccl5* from the endothelium in *Mrg15*^fl/fl^ mice (n=6) and *Mrg15*^iECKO^ mice (n=6). (L) Real time-qPCR analysis of the expression of *Mrg15*, *Itga5*, *Icam1*, *Vcam1*, *Ccl5* from the endothelium in *Mrg15*^WT^ mice (n=6) and *Mrg15*^iECOE^ mice (n=6). The statistical analysis was performed by two-tailed Student t test for J, *Mrg15*, *Itga5*, *Icam1* and *Ccl5* in K, and *Ccl5* in L, by Welch’s t test for I, *Ccl5* in K, and *Mrg15*, *Itga5*, *Icam1* in L, by Mann-Whitney test for *Vcam1* in L.

We then performed unbiased clustering analysis on the EC, which identified five distinct subpopulations, namely inflammatory EC1-EC3, extracellular matrix-associated EC, and lipid-handling EC (Figure 4F). We found that the proportion of inflammatory EC1 was significantly increased in the *Mrg15*^iECKO^ group (Figure 4F). Further pseudotime analysis revealed that *Mrg15* knockout promoted the transition of inflammatory EC2 and EC3, as well as extracellular matrix-related EC, toward inflammatory EC1 (Figure 4G). Additionally, *Mrg15* knockout enhanced *Icam1* expression specifically in inflammatory EC1 (Figure 4H). Further immunofluorescence staining showed that endothelial-specific knockout of *Mrg15* increased ICAM1 expression in endothelial cells within carotid atherosclerotic plaques (Figure 4I), whereas endothelial-specific overexpression of *Mrg15* reduced it (Figure 4J). Endothelial-specific deletion of *Mrg15* enhanced endothelial inflammation in the partial carotid ligation model, while endothelial-specific overexpression of *Mrg15* attenuated it, as demonstrated by real-time qPCR (Figure 4K and Figure 4L). These findings demonstrate that endothelial *Mrg15* deficiency fuels inflammatory signaling and pro-atherogenic responses.

### MRG15 Decreases ICAM and ITGA5 Expression and Monocyte Adhesion to ECs Induced by Disturbed Flow

To validate the single-cell sequencing results, we investigated the role of endothelial MRG15 in endothelial inflammation. Consistent with the single-cell sequencing results, we observed upregulated expression of *Icam1* and integrin subunit alpha 5 (*Itga5*) in primary mouse endothelial cells (MECs) isolated from *Mrg15*^iECKO^ mice compared with those from *Mrg15*^fl/fl^ mice (Figure 5A). Conversely, opposite effects were observed in MECs from *Mrg15*^iECOE^ mice, with significant downregulation of *Icam1* and *Itga5* expression (Figure 5B). We subsequently examined whether MRG15 contributes to the proinflammatory responses in endothelial cells under disturbed flow. OSS significantly upregulated the expression of ICAM1 and ITGA5 at both protein and mRNA levels in HUVECs. Knockdown of MRG15 further enhanced the effects of OSS stimulation on ICAM1 and ITGA5 expression (Figure 5C and Figure 5D). Furthermore, MRG15 knockdown promoted OSS-induced adhesion of monocytes to HUVECs (Figure 5E). Overexpression of MRG15 significantly attenuated the activating effects of OSS on endothelial cells (Figure 5F through Figure 5H). Meanwhile, *Mrg15* deletion potentiated TNF-α (tumor necrosis factor-α) -induced monocyte adhesion to MECs from *Mrg15*^iECKO^ mice relative to *Mrg15*^fl/fl^ controls (Figure 5I). These in vitro findings are highly consistent with the single-cell transcriptomic data and in vivo observations, demonstrating that MRG15 acts as a negative regulator in endothelial cell inflammation.

**Fig. 5.**
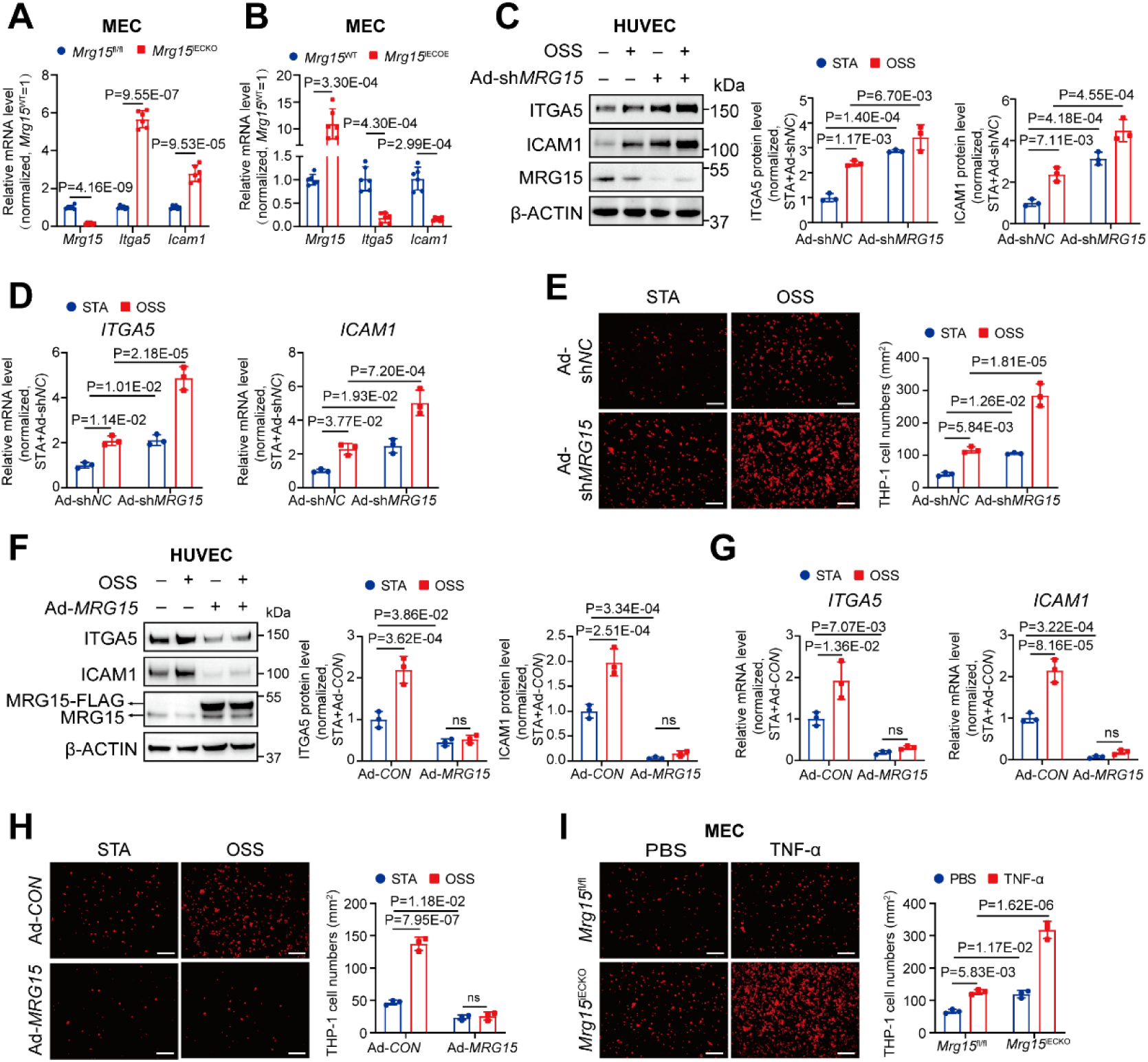
**MRG15 Decreases ICAM and ITGA5 Expression and Monocyte Adhesion to ECs Induced by Disturbed Flow** (A) Real time-qPCR analysis of the expression of *Mrg15, Itga5, and Icam1* in primary mouse endothelial cells (MECs) of *Mrg15*^fl/fl^ mice (n=6) and *Mrg15*^iECKO^ mice (n=6). (B) RT-qPCR analysis of the expression of *Mrg15, Itga5, and Icam1* in MECs of *Mrg15*^WT^ mice (n=6) and *Mrg15*^iECOE^ mice (n=6). (C) Western blot analysis of MRG15, ITGA5, ICAM1, and β-ACTIN levels in human umbilical vein endothelial cells (HUVECs) transfected with control shRNA or *MRG15* shRNA adenoviruses for 48 hours and then subjected to STA or oscillatory shear stress (OSS) (6 ± 0.5 dyn/cm^2^; 1 Hz; n=3) for 12 hours (n=3). (D) Real time-qPCR analysis of *ITGA5* and *ICAM1* levels in HUVECs transfected with control shRNA or *MRG15* shRNA adenoviruses for 48 hours and then subjected to STA or OSS (6 ± 0.5 dyn/cm^2^; 1 Hz; n=3) for 12 hours (n=3). (E) Representative images of monocyte-endothelial adhesion. HUVECs transfected with control shRNA or *MRG15* shRNA adenoviruses for 48 hours and then subjected to STA or OSS (6 ± 0.5 dyn/cm^2^; 1 Hz; n=3) for 12 hours (n=3) (Scale bar, 50 µm). (F) Western blot analysis of MRG15, ITGA5, ICAM1, and β-ACTIN levels in HUVECs transfected with Ad-*GFP* or Ad-*MRG15* for 48 hours and then subjected to STA or OSS (6 ± 0.5 dyn/cm^2^; 1 Hz; n=3) for 12 hours (n=3). (G) Real time-qPCR analysis of *ITGA5* and *ICAM1* levels in HUVECs transfected with Ad-*GFP* or Ad-*MRG15* for 48 hours and then subjected to STA or OSS (6 ± 0.5 dyn/cm^2^; 1 Hz; n=3) for 12 hours (n=3). (H) Representative images of monocyte-endothelial adhesion. HUVECs transfected with Ad-*GFP* or Ad-*MRG15* for 48 hours and then subjected to STA or OSS (6 ± 0.5 dyn/cm^2^; 1 Hz; n=3) for 12 hours (n=3) (Scale bar, 50 µm). (I) Representative images of monocyte-endothelial adhesion. Endothelial cells were isolated from *Mrg15*^fl/fl^ mice and *Mrg15*^iECKO^ mice, treated with PBS or human TNF-α (tumor necrosis factor-α; 10 ng/mL) for 24 hours (n=3) (Scale bar, 50 µm). The statistical analysis was performed by two-tailed Student t test for *Mrg15* in A, by Welch’s t test for *Itga5* and *Icam1* in A, and B, by two-way ANOVA followed by Tukey’s post test for C, D, E, F, G, H, and I.

### MRG15 recruits EZH2 to repress transcription of ICAM1 and ITGA5

To further elucidate the mechanisms by which MRG15 regulates ICAM1 and ITGA5 expression, we next examined whether ITGA5 and ICAM1 are transcriptional targets of MRG15. Cleavage Under Targets and Tagmentation (CUT&Tag) analysis demonstrated significant enrichment of MRG15 at the promoter regions of *ICAM1* and *ITGA5* in HUVECs (Figure 6A), suggesting transcriptional regulation of these genes. To determine whether MRG15 modulates *ITGA5* and *ICAM1* expression via chromatin remodeling, we performed Assay for Transposase-Accessible Chromatin with sequencing (ATAC-seq) on endothelial cells transduced with control or *MRG15* shRNA adenovirus. ATAC-seq revealed increased chromatin accessibility at the promoters of *ITGA5* and *ICAM1* upon MRG15 knockdown in HUVECs (Figure 6A).

**Fig. 6.**
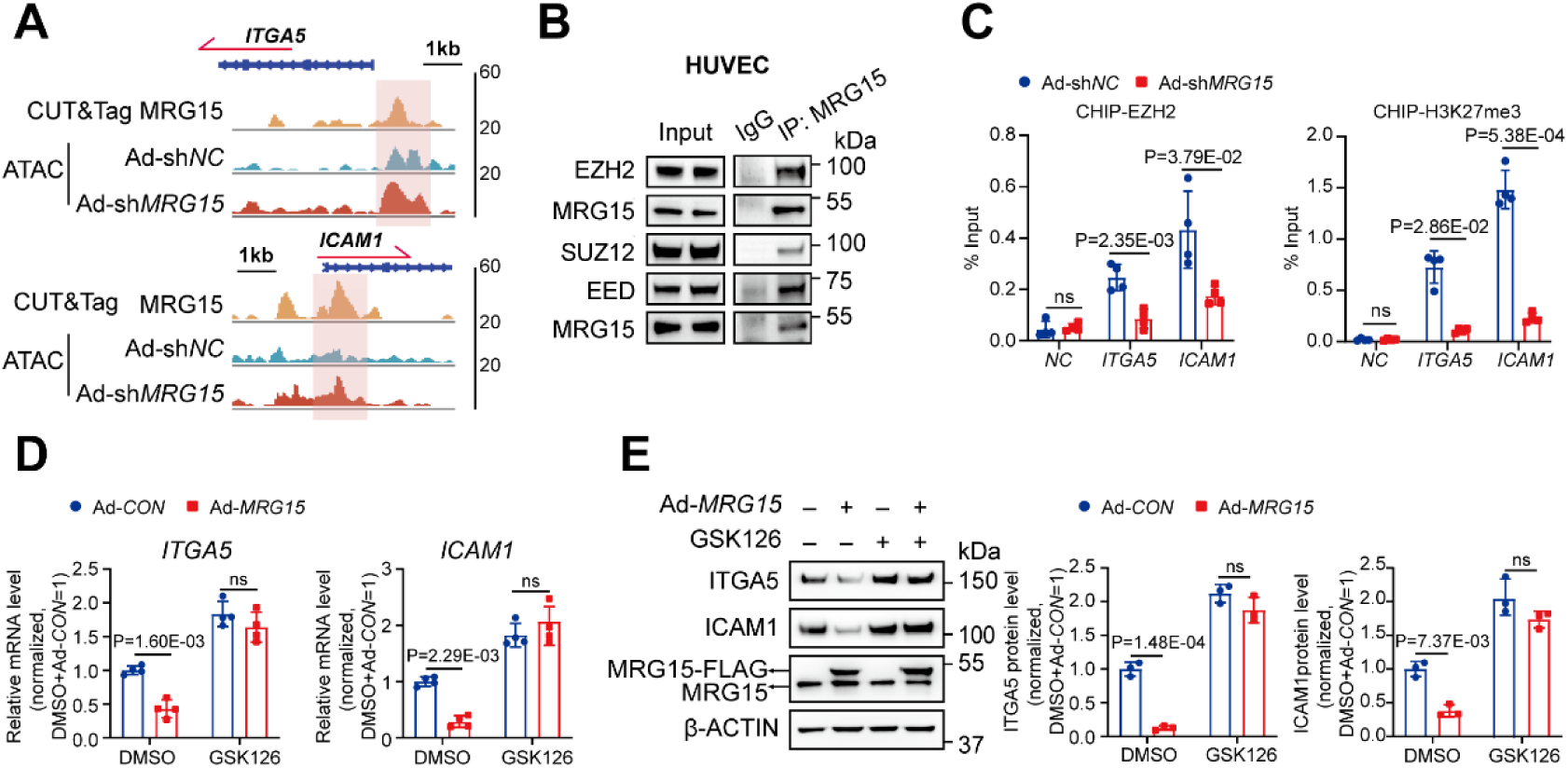
**MRG15 recruits PRC2 to repress transcription of ICAM1 and ITGA5** (A) Representative snapshots of Cleavage Under Targets and Tagmentation (CUT&Tag) tracks for MRG15 in human umbilical vein endothelial cells (HUVECs) and assay for transposase-accessible chromatin using sequencing (ATAC-seq) tracks at the promoters of *ITGA5* and *ICAM1* in HUVECs transfected with control shRNA or *MRG15* shRNA adenoviruses for 48 hours. (B) Coimmunoprecipitation of MRG15, EZH2, SUZ12, and EED in HUVECs. (C) left: chromatin immunoprecipitation followed by qPCR (ChIP-qPCR) analysis validated the enrichment of EZH2 at the promoters of *ITGA5* and *ICAM1* in HUVECs transfected with control shRNA or *MRG15* shRNA adenoviruses for 48 hours (n = 4). right: ChIP-qPCR analysis validated the enrichment of H3K27me3 at the promoters of *ITGA5* and *ICAM1* in HUVECs transfected with control shRNA or *MRG15* shRNA adenoviruses for 48 hours (n = 4). (D) Real time-qPCR analysis of *ITGA5* and *ICAM1* levels. HUVECs were transfected with Ad-*GFP* or Ad-*MRG15* for 48 hours and then exposed to DMSO or EZH2 inhibitor GSK126 (1 μM) for 24 hours (n=3). (E) Western blot analysis of MRG15, ITGA5, ICAM1, and β-ACTIN levels. HUVECs were transfected with Ad-*GFP* or Ad-*MRG15* for 48 hours and then exposed to DMSO or EZH2 inhibitor GSK126 (1 μM) for 24 hours (n=3). The statistical analysis was performed by two-way ANOVA followed by Tukey’s post test for D and E, by Mann-Whitney test for NC in CHIP-EZH2 and ITGA5 in CHIP-H3K27me3 in C, by two-tailed Student t test for ITGA5 in CHIP-EZH2 and NC in CHIP-H3K27me3 in C, by Welch’s t test for ICAM1 in CHIP-EZH2 and ICAM1 in CHIP-H3K27me3 in C .

Previous studies have shown that MRG15 interacts with enhancer of zeste homolog 2 (EZH2) to regulate pluripotency in stem cells(*45*). Endogenous co-immunoprecipitation in HUVECs confirmed that MRG15 physically associates with EZH2, as well as with the other polycomb repressive complex 2 (PRC2) subunits, suppressor of zeste 12 protein homolog (SUZ12), and embryonic ectoderm development (EED) (Figure 6B). To determine whether MRG15 recruits EZH2 to the *ICAM1* and *ITGA5* promoters, we performed chromatin immunoprecipitation followed by qPCR (ChIP-qPCR) for EZH2 and the repressive histone mark H3K27me3 in control and MRG15-knockout HUVECs. MRG15 depletion markedly reduced the enrichment of both EZH2 and H3K27me3 at these promoters (Figure 6C), consistent with the observed increase in chromatin accessibility. Furthermore, treatment with the EZH2 inhibitor GSK126 attenuated the suppression of ICAM1 and ITGA5 expression induced by MRG15 overexpression at both mRNA and protein levels (Figure 6D and Figure 6E). Collectively, these results demonstrate that MRG15 recruits EZH2 to maintain repressive H3K27me3 marks at the *ICAM1* and *ITGA5* promoters, thereby suppressing their expression in endothelial cells.

## Discussion

Large-scale genome-wide association studies have identified multiple coronary artery disease risk loci at or near the MRG15 locus(*29–31*), yet the functional contribution of MRG15 to atherosclerosis has remained largely elusive. In the present study, we provided evidence that MRG15 acted as a critical negative regulator of endothelial activation in response to OSS. OSS stimulation rapidly induced neddylation in endothelial cells, leading to accelerated degradation of MRG15. Endothelial-specific deletion of MRG15 exacerbated atherosclerotic lesion formation in both partial carotid ligation and chronic high-fat diet-fed mouse models, whereas MRG15 overexpression in endothelial cells conferred protection against plaque development. Mechanistically, MRG15 recruited EZH2 to to maintain repressive H3K27me3 marks on the promoters of *ICAM1* and *ITGA5*.

Despite the widespread use of statins as first-line lipid-lowering therapy, a substantial proportion of patients on long-term treatment continue to develop or progress atherosclerotic disease. Therefore, beyond lipid-lowering therapies, the development of anti-inflammatory treatments holds significant importance for improving the prognosis of patients with atherosclerosis. Disturbed flow is a critical biomechanical stimulus that drives endothelial cells toward a proinflammatory phenotype(*8*). In the present study, we integrated CAD GWAS data with downregulated proteins in proteomic profiles from HUVECs under disturbed flow, demonstrating that MRG15 acts as a key endothelial mechanosensor in response to shear stress, thereby suppressing disturbed flow-induced atherosclerosis. Our results demonstrate that OSS stimulation downregulates MRG15 expression, thereby activating endothelial inflammation and accelerating atherosclerosis progression, whereas endothelial-specific overexpression of MRG15 effectively inhibits atherosclerosis. This is primarily achieved by driving an endothelial phenotypic switch from low to high *Icam1* expression and by activating integrin-mediated pathways in partial carotid ligation mouse models. These findings reveal that endothelial MRG15 represents a novel anti-inflammatory target for the treatment of atherosclerosis.

Neddylation is an important ubiquitin-like post-translational modification that plays a critical role in regulating endothelial cell function. Recent studies have shown that neddylation inhibitors can attenuate atherosclerotic plaque formation in mice(*46*). However, whether neddylation is involved in disturbed flow-induced endothelial inflammatory activation remains unclear. Previous studies have reported that MRG15 is one of the proteins most significantly upregulated following treatment with neddylation inhibitors(*47*). Nevertheless, whether neddylation directly controls MRG15 degradation is still unknown. We found that disturbed flow rapidly induced the level of neddylation in endothelial cells, resulting in neddylation-dependent degradation of the MRG15 protein. Loss of MRG15 stability under OSS, thereby amplifying endothelial proinflammatory responses and accelerating atherogenesis. Our results demonstrate that neddylation represents a novel post-translational modification in endothelial cells that responds to blood flow shear stress. It is rapidly activated under disturbed flow conditions, with the MRG15 serving as a key target protein of neddylation.

EZH2 serves as the catalytic subunit of Polycomb repressive complex 2 (PRC2), which catalyzes trimethylation of histone H3 at lysine 27 (H3K27me3) to mediate transcriptional repression(*48*). Knockdown of EZH2 in endothelial cells upregulates cell adhesion-related pathways(*49*). Notably, EZH2 binds to the YAP promoter and represses its expression(*50*), whereas YAP serves as a key proinflammatory factor in endothelial cells in response to disturbed blood flow. However, previous studies have shown that EZH2 expression is upregulated under OSS(*51*). These findings suggest that mechanisms beyond alterations in EZH2 expression regulate its recruitment to distinct promoters under OSS. ChIP-seq analysis of endothelial cells treated with STA or OSS revealed inconsistent EZH2 enrichment patterns across different gene promoter regions(*52*). Our findings provide new insights into the regulation of EZH2 activity in endothelial cells under shear stress: MRG15 recruited EZH2 to maintain repressive H3K27me3 marks at the *ICAM1* and *ITGA5* promoters. Under OSS, reduced MRG15 levels lead to diminished EZH2 and H3K27me3 occupancy at these loci, thereby derepressing *ICAM1* and *ITGA5* transcription, activating endothelial inflammation, and promoting atherogenesis.

Collectively, our findings identify MRG15 as a a mechanosensitive suppressor of atherosclerosis induced by disturbed flow. Disturbed blood flow rapidly led to an elevation of protein neddylation, which subsequently induced neddylation-dependent degradation of MRG15 within endothelial cells. This degradation of MRG15 alleviated the transcriptional repression of ICAM1 and ITGA5, thereby enhancing monocyte adhesion to ECs. These results position MRG15 emerges as a promising novel therapeutic target for atherosclerosis.

